# Phosphate and osmotic adaptation: a major role for phosphate in charge balance and metabolic responses in *Escherichia coli*

**DOI:** 10.64898/2026.06.06.730615

**Authors:** Debbie McLaggan, Wolfgang Epstein

## Abstract

Adaptation of Escherichia coli to osmotic upshift requires rapid accumulation of intracellular solutes to restore turgor and maintain cellular homeostasis. While compatible solutes are well-established contributors to this process, they do not fully account for the early events following osmotic stress. Here, we demonstrate that inorganic phosphate and phosphorylated metabolites play a major and previously underappreciated role in osmoadaptation.

Following osmotic upshift under conditions where accumulation of compatible solutes is restricted, E. coli exhibits a substantial increase in intracellular phosphate after a short lag. This increase accounts for a significant fraction of the charge balance required during rapid uptake of K^+^ and NH_4_^+^, the latter supporting glutamate synthesis as a principal counterion. Concomitantly, nucleotide pools display complex, multiphasic dynamics, including a transient decrease in adenylate energy charge whose duration correlates with stress magnitude. In addition, levels of pyrophosphate and key glycolytic intermediates, including dihydroxyacetone phosphate and 1,3-bisphosphoglycerate, increase markedly, indicating redistribution of phosphate into central metabolic pathways.

These findings support a model in which phosphate uptake and metabolic redistribution contribute both to intracellular charge balance and to dynamic metabolic reorganisation during osmotic stress. By linking ion transport with central metabolism, this work expands current models of bacterial osmoadaptation and identifies phosphate flux as a key component of the early stress response.

**IMPORTANCE:** Bacterial survival in fluctuating environments depends on rapid adaptation to osmotic stress. While compatible solutes are central to this process, their contribution does not fully account for early events in *Escherichia coli* following osmotic upshift. This work demonstrates that inorganic phosphate uptake and redistribution into nucleotide and glycolytic pools contribute substantially to balance the large positive charge entering the cell as it takes up K^+^ and NH_4_^+^ during osmotic upshift. These findings expand current models of bacterial osmoregulation by identifying phosphate flux as a central integrator of ion homeostasis and metabolic adaptation.

## INTRODUCTION

Bacteria must continuously adapt to changes in environmental osmolarity to maintain cellular integrity and physiological function (1, 2). In *Escherichia coli*, increases in external osmolarity (osmotic upshift) trigger rapid adjustments in intracellular solute composition that restore turgor pressure and preserve macromolecular activity. The early phase of osmoadaptation is characterised by rapid uptake of K^+^ through multiple transport systems, followed by accumulation of glutamate, which serves as the principal counterion (3–7). Subsequently, osmolytes such as trehalose and glycine betaine accumulate and progressively replace K^+^, thereby stabilising proteins and cellular structures without interfering with metabolism (8–10).

These core features of osmoadaptation are well established; however, increasing evidence suggests that they represent only part of a broader, highly integrated physiological response. Recent studies emphasise that ion transport, metabolic flux, and regulatory signalling networks are tightly coordinated during stress adaptation, rather than operating as independent processes (11-14). In particular, potassium transport systems are now understood to exhibit complex regulation and functional specialisation, responding dynamically to environmental and metabolic cues (5,13). This emerging perspective highlights the need to consider osmoadaptation within the context of whole-cell physiology.

While osmolytes and ion fluxes have been extensively characterised, comparatively little attention has been paid to the role of phosphate and phosphorylated metabolites during osmotic stress. Phosphate is central to cellular metabolism, participating in energy transfer, nucleotide synthesis, and regulation of metabolic pathways. Early studies reported changes in adenine nucleotide pools following osmotic shifts (15), suggesting that osmotic stress perturbs cellular energy balance. The adenylate energy charge, a key indicator of metabolic state, is known to respond sensitively to environmental and physiological perturbations (16). More recent work has extended these observations, demonstrating that environmental stresses induce coordinated changes across metabolic networks, including central carbon metabolism and nucleotide biosynthesis (11, 17,18). In parallel, global regulatory mechanisms such as the stringent response play a central role in integrating metabolic and environmental signals. The alarmone (p)ppGpp modulates transcription, translation, and metabolic fluxes in response to stress, linking nutrient availability with growth control (12,). Activation of the stringent response during osmotic stress further underscores the extent to which osmoadaptation involves coordinated global regulation rather than simple accumulation of osmolytes.

Despite these advances, the potential contribution of phosphate fluxes to osmotic adaptation has not been systematically examined. In particular, the role of inorganic phosphate in balancing the large influx of cations during early osmoadaptation, and its redistribution into intracellular metabolic pools, remain poorly understood. Earlier work demonstrated links between potassium and phosphate transport systems (19, 20), but the physiological significance of this relationship in the context of osmotic stress has not been fully resolved.

Here, using radiolabeling, biochemical assays, and nucleotide profiling, we investigate the role of phosphate during osmotic upshift in *E. coli*. We show that osmotic stress leads to a substantial increase in intracellular phosphate, which is redistributed into nucleotide pools and glycolytic intermediates. These changes are associated with dynamic alterations in energy charge and metabolite levels, consistent with a coordinated metabolic response. Our findings indicate that phosphate uptake and metabolism contribute significantly to intracellular charge balance during cation influx and form an integral component of osmoadaptation. This work expands current models by identifying phosphate as a key link between ion homeostasis and metabolic regulation during bacterial stress response. By integrating ion transport with metabolic reorganisation, this study provides a revised framework for understanding osmoadaptation as a coordinated physicochemical and metabolic process.

## RESULTS

### Phosphate is required for maximal K^+^ uptake following upshift

K^+^ uptake following osmotic upshift was reduced both in initial rate and extent in phosphate limited medium. (Fig. 1). However, this reduction in initial uptake rate was dependent on the presence of NH_4_^+^; in the absence of both phosphate and nitrogen, the initial rate of K^+^ uptake was restored although total accumulation remained lower. This effect was independent of external pH; similar results were obtained at pH 7.0 and 8.2 (data not shown). Measurements of ^86^Rb^+^ efflux (K^+^ tracer) indicated that reduced net uptake was not due to increased efflux, demonstrating that phosphate limitation specifically affects K^+^ influx under these conditions.

**FIG 1.**
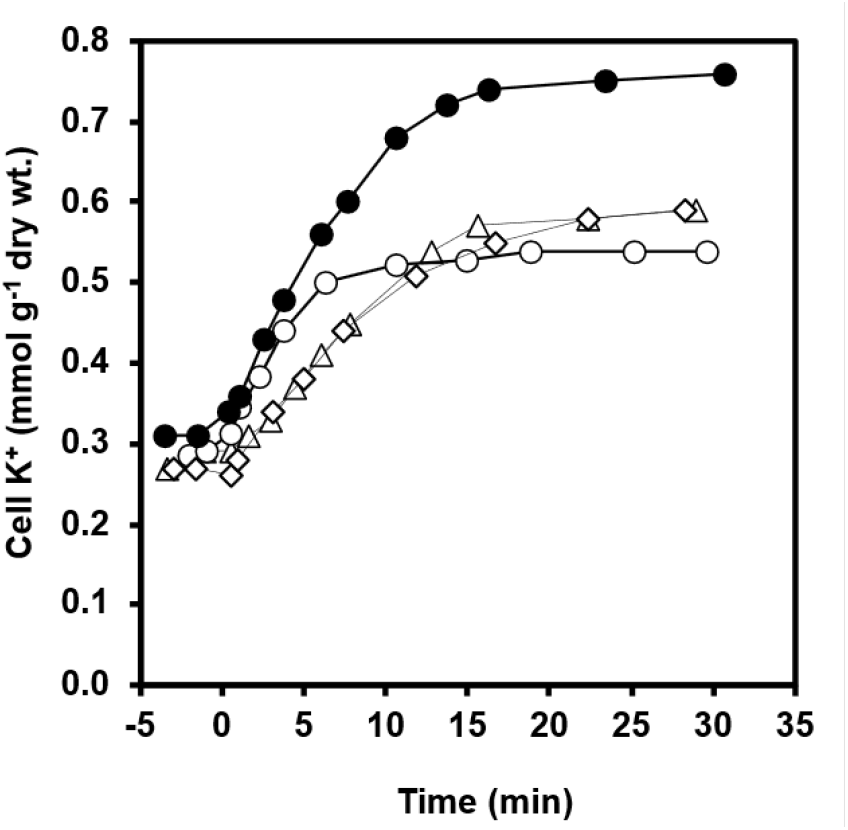
Phosphate and nitrogen dependence of K^+^ uptake following osmotic upshift. Cells in exponential phase were transferred to complete medium with 10 mM Na_2_HPO_4_ and 8 mM NH_4_SO_4_ (•), medium lacking phosphate (△, ⋄), or medium lacking both phosphate and nitrogen (⋄), and subjected to moderate osmotic upshift 0.57 M NaCl (•,△, ◯) or 0.9 M glucose (⋄) at t = 0.

### Osmotic upshift increases intracellular phosphate content

Osmotic upshift led to a reproducible increase in total intracellular phosphate (Fig. 2). Following a lag of approximately 2 min, phosphate levels increased by ∼30% relative to control cells. This increase in phosphate was dependent on K^+^ availability; and was not observed under K^+^-limiting conditions.

**FIG 2.**
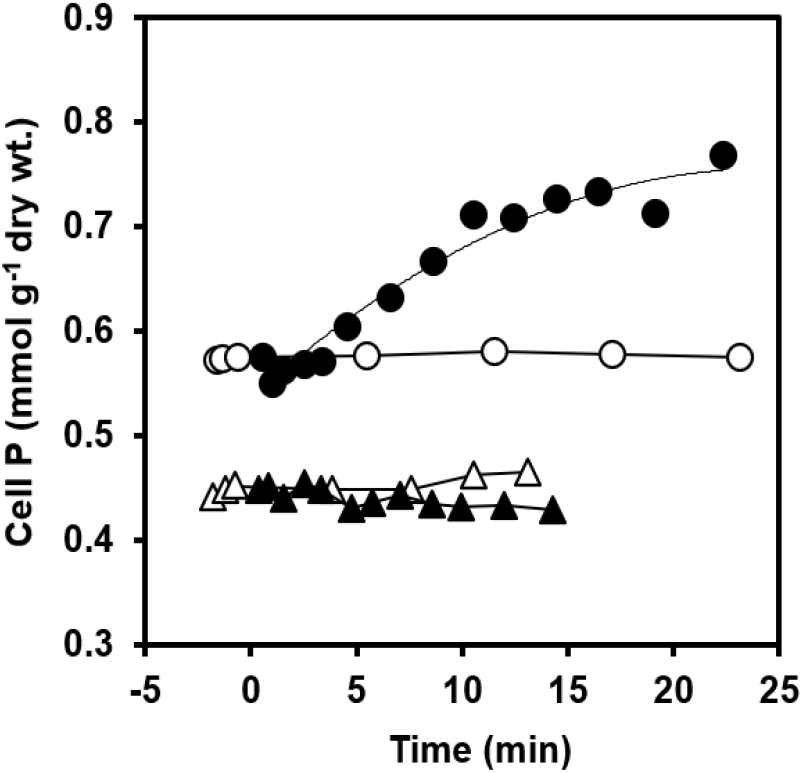
Increase in intracellular phosphate following osmotic upshift. Cells grown in medium containing ^32^P (0.3 mM; 1.67 Ci/mmol) with excess K+ (5 mM, ◯,•) or limiting K^+^ (0.3 mM, △▴) were subjected to moderate osmotic upshift at t = 0 (0.9 M glucose (▴,•). Intracellular phosphate exhibited a lag of approximately 2 min followed by an increase of ∼30%.

The magnitude of phosphate accumulation corresponded to approximately 25% of the K^+^ accumulated over the same period, indicating that phosphate contributes substantially to intracellular charge balance during early osmoadaptation.

### Glycolytic intermediates accumulate transiently following upshift

Among glycolytic intermediates assayed, only dihydroxyacetone phosphate and 1,3-bisphosphoglycerate showed substantial increases following osmotic upshift (Fig. 3). These increases were transient after moderate upshift conditions but persisted longer in the absence of a nitrogen source (Fig. 3), suggesting that synthesis of glutamate or other N-containing compounds facilitates the return to control levels. The magnitude of phosphate accumulation in these intermediates accounted for a substantial fraction of total phosphate uptake, suggesting that central metabolism could act as a reservoir for excess phosphate during osmotic stress.

**FIG 3.**
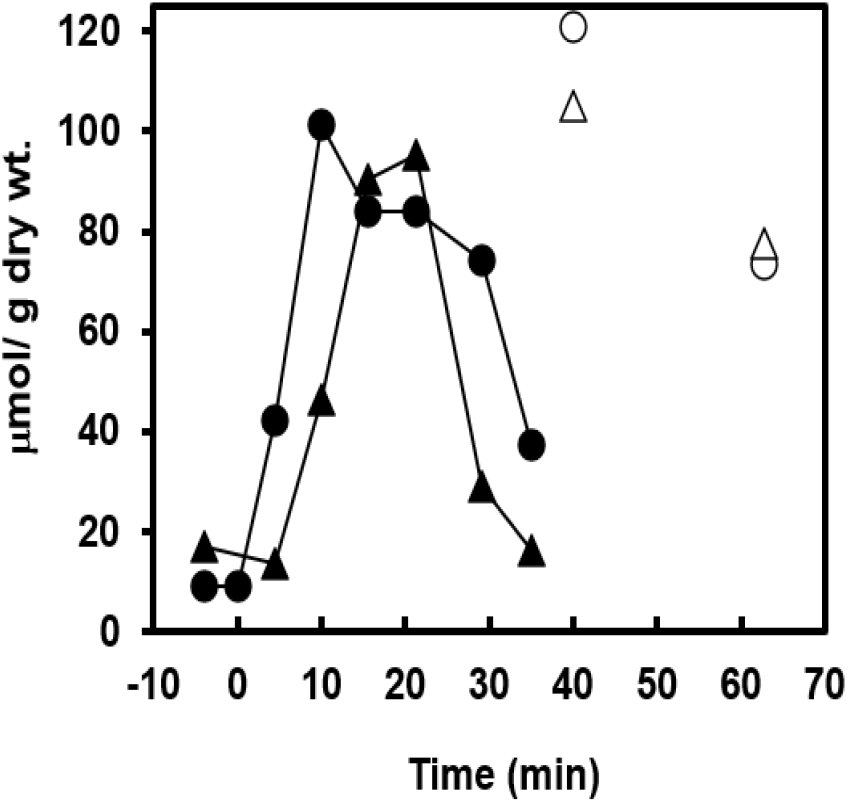
Accumulation of glycolytic intermediates following osmotic upshift. Changes in dihydroxyacetone phosphate (DHAP,•, ◯), and 1,3-bisphosphoglycerate (1,3-BPG, ▴△), following moderate osmotic upshift (0.9 M glucose) are shown in the presence (closed symbols) or absence (open symbols) of a nitrogen source. Increases were transient in the presence of nitrogen but prolonged under nitrogen limitation.

### Response of nucleotide pools to osmotic upshift

Adenine nucleotide pools displayed multiphasic changes (Fig. 4). After upshift there was an immediate fall in ATP and immediate rises in ADP and AMP. After mild or moderate upshock this was followed within a few minutes by a large increase in ATP and reductions in ADP and AMP levels. The increase in ATP levels was of relatively short duration after mild upshock (Fig. 4) but a high level of ATP persisted for several hours after moderate upshock (Fig. 5A), well into the period where the cells had resumed exponential growth. After severe upshift the phase of low ATP was prolonged whereas the elevated ADP and AMP returned to control levels within an hour (Fig.5B).

**FIG 4.**
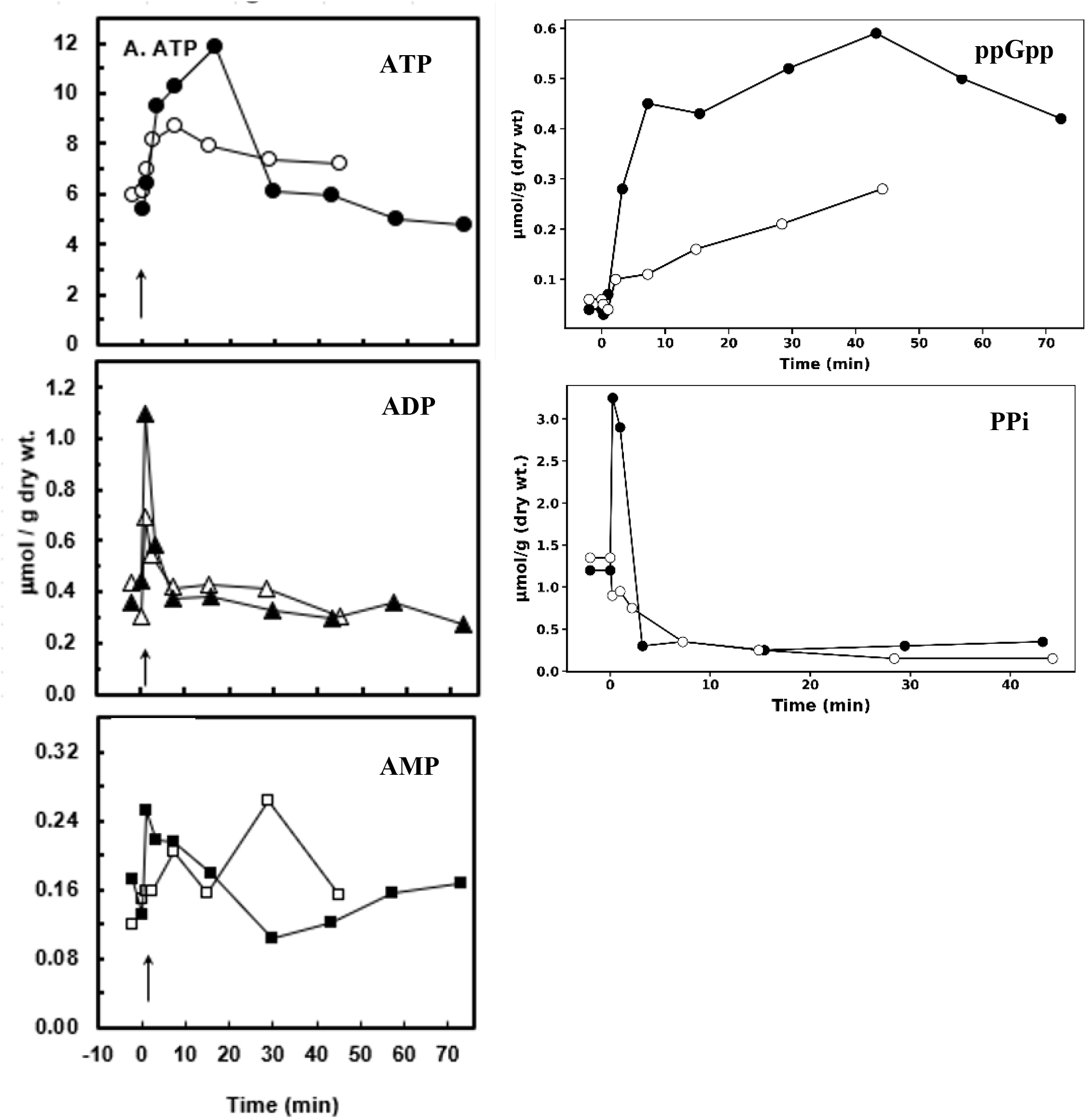
Nucleotide pools following mild osmotic upshift. Cells were subjected to mild osmotic upshift (0.45 M NaCl) at t = 0. Intracellular nucleotide levels were determined over time by radiolabeling and thin-layer chromatography. Closed symbols excess K^+^ (5mM); open symbols limiting K^+^ (0.3 mM).

**FIG 5.**
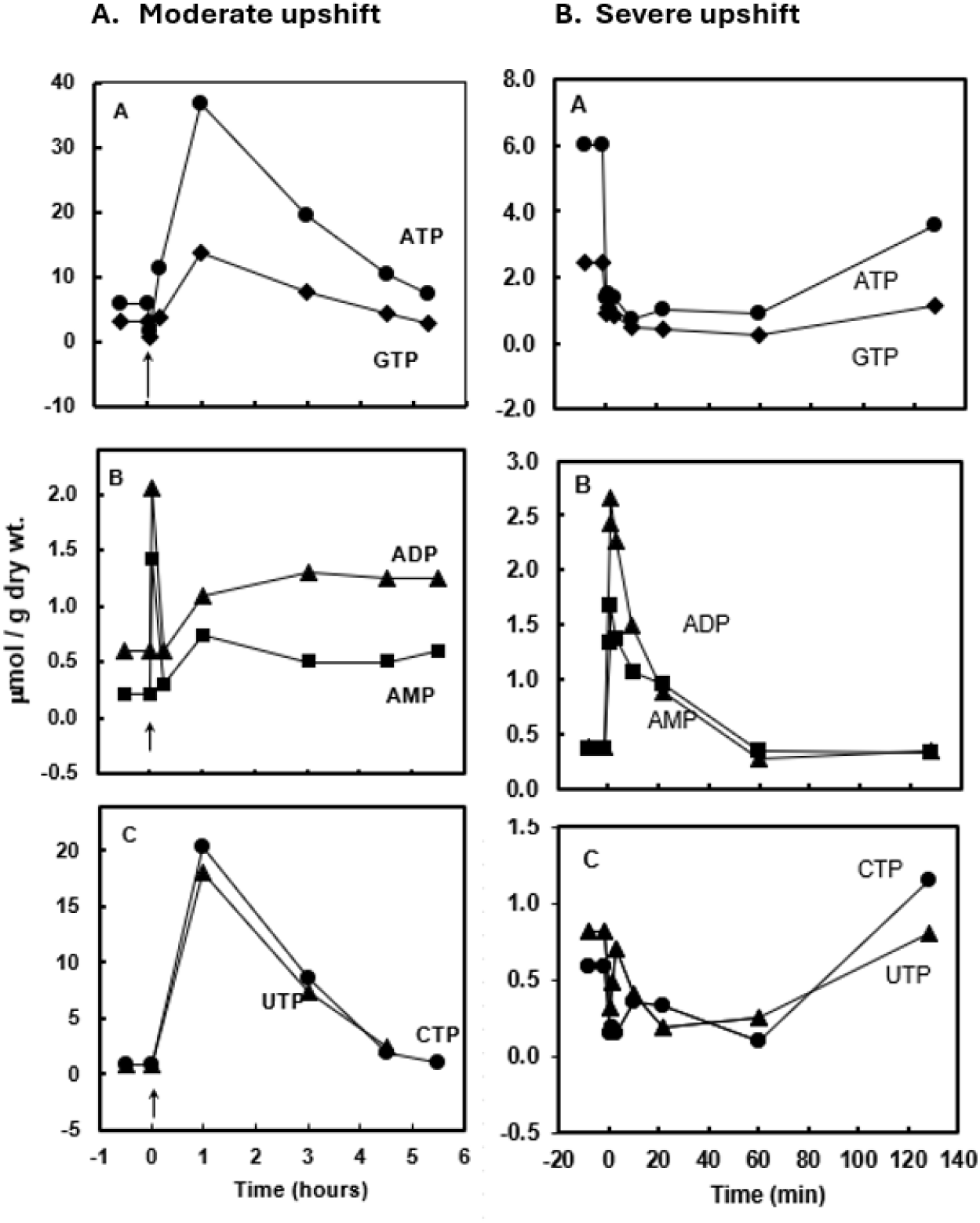
Nucleotide pools following moderate and severe osmotic upshift. Changes in nucleotide pools following moderate osmotic upshift (0.57 M NaCl) are shown in panel A and following severe osmotic upshift (0.65M NaCl) in panel B. . Assumed control value for ATP, 6 μmol g^-1^ dry wt. Nucleotide concentrations following severe osmotic upshift are shown. Prolonged alterations in ATP, ADP, and AMP levels were observed, reflecting sustained metabolic perturbation

The changes in adenine nucleotide levels resulted in a reduction in adenylate energy charge (Fig. 6). The duration of the low-energy state correlated with stress severity, lasting minutes under mild upshift conditions but over an hour under severe osmotic stress (Fig. 5B).This is consistent with the known sensitivity of the adenylate energy charge to changes in cellular metabolic state (12).

**FIG 6.**
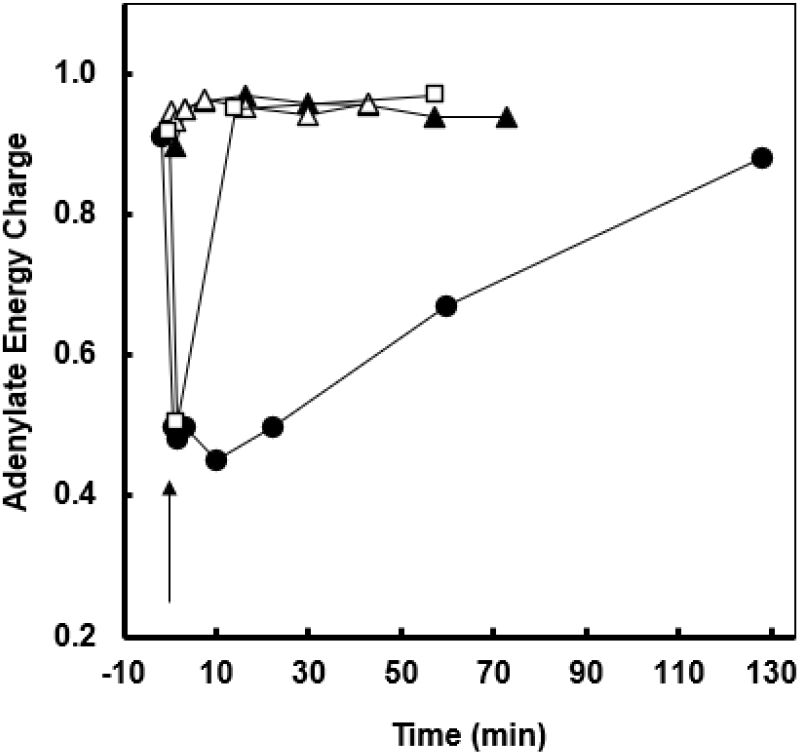
Changes in adenylate energy charge following osmotic upshift. Adenylate energy charge was calculated from ATP, ADP, and AMP levels following mild moderate, and severe osmotic upshift. (▴,0.45M NaCl excess K^+^) (□ 0.57M NaCl excess K^+^) (• 0.65M NaCl excess K^+^) (△0.45M NaCl limited K^+^).

Other nucleotide pools showed similar trends, with GTP, dATP, and dGTP paralleling ATP dynamics (supplementary Fig.S1) and CTP and UTP exhibiting larger and more sustained increases (Fig. 5).

Levels of ppGpp rose moderately after mild upshift but reached peak values that were more than 10 times those of the control after moderate upshift (Fig. 5A). Levels of ppGpp remained high to the end of the time of sampling as would be expected given the correlation of high ppGpp levels with slow growth rate (21). pppGpp also transiently rose 4-fold after mild upshock (data not shown). Pyrophosphate pools exhibited a diphasic response to mild upshift, more than doubling in the first minute to falling to levels lower than pre-upshift (Fig. 4). This rapid decrease in PPi was confirmed chemically (22).

### Effect of limiting K+ on nucleotide pools following mild upshock

When K^+^ uptake was prevented, most of the subsequent events were much reduced or absent, changes in most purine nucleotides were much less marked, ppGpp increased at a slower rate and the initial transient rise in PPi was not observed (Fig. 4).

### Effect of upshock on nucleotides in the presence of chloramphenicol

Inhibition of protein synthesis resulted in similar changes. After an early drop in all nucleotides measured, by 15 min they have risen to levels that are similar to those achieved in the absence of chloramphenicol PPi levels were also observed to drop (data not shown).

Energy charge decreased transiently, with the duration of the decrease correlating with stress magnitude.

## DISCUSSION

The results presented here demonstrate that phosphate uptake and redistribution into metabolic pools make a substantial contribution to the early response to osmotic upshift in *Escherichia coli*. Classical models of osmoadaptation emphasise K^+^ uptake and osmolyte accumulation as primary determinants of turgor restoration (1, 2), but more recent work highlights the importance of integrating ion transport with broader metabolic and regulatory networks (13, 14). Within this framework, our findings identify phosphate metabolism as a key and previously underappreciated component of the osmotic stress response.

The requirement for phosphate in supporting K^+^ uptake under conditions where NH_4_^+^ is present suggests a role in maintaining electroneutrality during rapid cation influx. The cell may not be able to maintain charge balance solely by proton extrusion during the early phase of osmotic upshift, when uptake of both K^+^ and NH_4_^+^ is high. In this context, uptake of divalent phosphate provides an efficient means of counterbalancing positive charge. This interpretation is consistent with earlier work linking potassium and phosphate transport (19,20, 21), but also aligns with more recent studies showing tight coupling between ion homeostasis, membrane energetics, and metabolic flux in bacteria (7, 14),).

An important feature of the response observed here is the redistribution of phosphate into nucleotide and glycolytic intermediate pools. We propose that once immediate biophysical requirements such as maintenance of intracellular pH and membrane potential are stabilised, excess phosphate is transiently stored in central metabolic intermediates. The accumulation of dihydroxyacetone phosphate and 1,3-bisphosphoglycerate supports the idea that central carbon metabolism acts as a dynamic reservoir for phosphate during stress. This interpretation is consistent with systems-level analyses showing that metabolic networks are rapidly reconfigured in response to environmental perturbations, with central metabolism playing a buffering and regulatory role (11). More recent metabolomics studies further emphasise that stress responses involve coordinated redistribution of metabolic intermediates rather than simple linear pathway activation, reinforcing the concept of metabolism as a flexible reservoir system during adaptation.

The complex, multiphasic responses of nucleotide pools provide further evidence for coordinated metabolic regulation. The transient decrease in adenylate energy charge reflects an immediate disturbance in energy balance following osmotic upshift, followed by recovery as adaptation proceeds. Such dynamics are characteristic of stress responses and are increasingly understood as part of global regulatory programs linking metabolism, growth, and environmental sensing. In particular, the pronounced accumulation of (p)ppGpp observed here is consistent with activation of the stringent response, which integrates nutrient status, stress signalling, and growth control (12).

Recent reviews have highlighted the central role of (p)ppGpp in reshaping transcriptional and metabolic networks under stress conditions, including osmotic and ionic stresses, further supporting the interpretation that osmoadaptation involves coordinated global regulation rather than isolated transport events

The prolonged perturbations observed under more severe osmotic conditions suggest that the duration of metabolic imbalance scales with stress magnitude. This is in agreement with recent studies demonstrating that bacterial responses to environmental stress are graded and involve dynamic transitions between physiological states, rather than simple on/off responses. In this context, the correlation between stress severity, adenylate energy charge, and nucleotide pool dynamics observed here provides a useful quantitative link between environmental perturbation and cellular metabolic state.

Taken together, these findings extend current models of osmoadaptation by incorporating phosphate metabolism as an integral component of the response. Rather than relying solely on accumulation of compatible solutes, *E. coli* employs a coordinated strategy involving ion transport, metabolic reorganisation, and redistribution of phosphorylated compounds. Within this framework, phosphate serves both as a charge-balancing species and as a central element of metabolic adaptation.

Although the experiments reported here were performed using classical biochemical approaches, the results are highly consistent with more recent systems-level and multi-omics analyses of bacterial stress responses (17, 18). These modern studies emphasise the integration of metabolic flux, ion transport, and regulatory networks as a general principle of bacterial adaptation. Our findings therefore provide a mechanistic underpinning for these broader observations and highlight phosphate flux as a key variable linking ion homeostasis with central metabolism.

In conclusion, phosphate is not simply a background metabolite but a major component of the osmotic stress response, contributing both to charge balance and to metabolic adaptation. Incorporating phosphate into models of osmoadaptation provides a more complete understanding of how bacterial cells coordinate physical and metabolic processes to survive fluctuating environments

## MATERIALS AND METHODS

### Growth conditions and osmotic upshift

*Escherichia coli* strain FRAG5 (**F**^***-***^ ***gal thi*** *rha lacz Δ* ***(kdpABC)5*)** and its *otsA* derivative FRAG69, were grown at 37° C in Na-HEPES-buffered minimal medium (0.17 osM) containing 0.1 M HEPES-Na (pH 7.4), 5 mM KCl, 10 mM Na_2_HPO_4_, 8 mM (NH_4_)_2_SO_4_, 1 mM sodium citrate, 0.4 mM MgSO_4_, 1 µg ml_−1_ thiamine, and 10 µM FeSO_4_, with 10 mM glucose as the carbon source (6). Unless otherwise indicated, experiments were performed using strain FRAG69. FRAG5 was used specifically for analysis of glycolytic intermediates.

Growth was monitored by turbidity measurements in 18 mm tubes at 610 nm (Bausch & Lomb Spectronic 20 colorimeter) and converted to dry weight values based on standard curves at low and at high osmolarity. Osmotic upshift was performed during exponential growth (250–350 µg dry weight ml_−1_) by addition of 2.5M NaCl or 2.7M glucose in growth medium to the indicated final osmolarities. For nucleotide analysis, upshift was with NaCl since glucose interfered with subsequent sample analysis. For glycolytic intermediate analysis, upshift was with glucose since NaCl inhibited some of the enzymatic assays.

Mild, moderate, and severe upshift conditions corresponded to final osmolarities of:

- Mild: 1.02 osM (0.45M NaCl)
- Moderate: osM, 1.24/1.26 osM (0.57M NaCl/0.9M glucose)
- Severe 1.39 osM (0.65 M NaCl)

### Measurement of intracellular phosphate and K^+^

Cells were grown for 4 generations in Na-Hepes medium. For intracellular phosphate determination the media phosphate concentration was reduced to 0.3 mM and ^32^P0_4_ at 1.67 Ci/mmol was added prior to upshift. Cells were collected on gridded 18 mm membrane filters (pore size 0.45 μm, Millipore type HA) and washed with hypertonic buffer. Intracellular phosphate was quantified using ^32^P labeling and scintillation counting. Potassium levels were measured by flame photometry (23).

### Efflux of ^86^Rb^+^

Cells were grown for 3 generations in Na-Hepes medium (K^+^ 1mM, ^86^Rb^+^ 5 nCi/ml, 0.02 mM). Prior to upshift the cells were collected on a 47 mm membrane filter, washed and suspended in Na-Hepes medium (K^+^ 10 mM).

### Glycolytic intermediates

Metabolites were extracted by cold osmotic shock as described (9), and quantified enzymatically by coupling reactions to NAD(P)H oxidation or reduction, monitored at 340 nm (24-26). The following were assayed: Glucose-6-P, glucose-1-P, fructose-6-P, fructose-1,6-bis-P, dihyroxyacetone phosphate (DHAP), glyceraldehyde-3-P, 1,3-bis-phosphogycerate (1,3-BPG), and 3-phosphoglycerate (3-PG), P-enolpyruvate, pyruvate, and 2-P-glycerate. All enzymes for the assays were from Sigma Chemical Co.

### Nucleotide analysis

Cells were grown in small volumes, 0.6 - 1.5 ml, in medium containing 0.3 mM phosphate and ^32^PO_4_ (100 Ci/mol) in 13 x 100 mm tubes that were placed inside 125 ml Erlenmeyer flasks containing 6 ml water to reduce evaporative loss. Growth was monitored by turbidity measurements of parallel cultures lacking radioactivity. Labeling was arrested by chilling on ice, addition of formic acid and precipitation of inorganic phosphate with tungstate, tetraethylammonium and procaine as described by Bochner and Ames 27). After centrifugation at 4° C, the supernatants were stored at -80° C until analysis. Nucleotides were separated by two-dimensional thin-layer chromatography as described by Bochner and Ames (27) and quantified by radiolabel incorporation, using ATP as an internal reference standard. Total phosphate content was measured on cell pellets obtained from 25 - 50 ml of cultures that were washed with 60% ethanol, dried to constant weight in vacuo and weighed. The pellets were digested with concentrated perchloric acid (60%) at 160° C until the samples were colorless, after which phosphate was assayed as described by Chen et al.(28). The weights of pellets from 25 - 50 ml cultures was used to convert turbidity measurements to dry weight per ml. Dry weight was assumed to remain constant for 20 min after upshift; after that it was corrected by any change in turbidity. All experiments were performed at least twice with similar results; representative data are shown.

## ACKNOWLEDGEMENTS

We thank Dan Fraenkel and Ian Booth for helpful comments. This work was supported in part by grant DCB-8704059 from the National Science Foundation and grant GM2323323 from the National Institutes of Health.

## AUTHOR CONTRIBUTIONS

## ADDITIONAL FILES

### Supplemental Material

Phosphate and osmotic adaptation Fig S1 and Fig S1 additional nucleotides

