## Supplemental Fig S1, Fig S2 for "Phosphate and osmotic adaptation: a major role for phosphate in charge balance and metabolic responses in *Escherichia coli*"

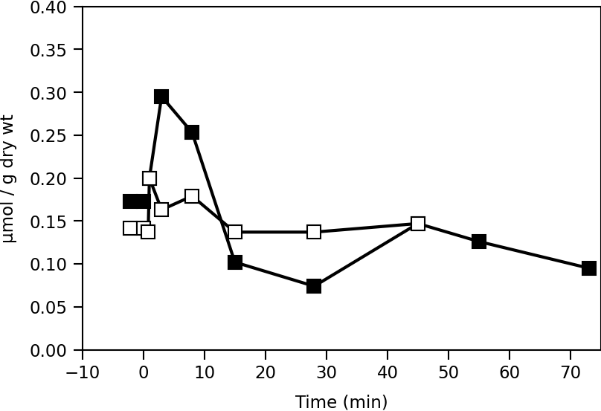

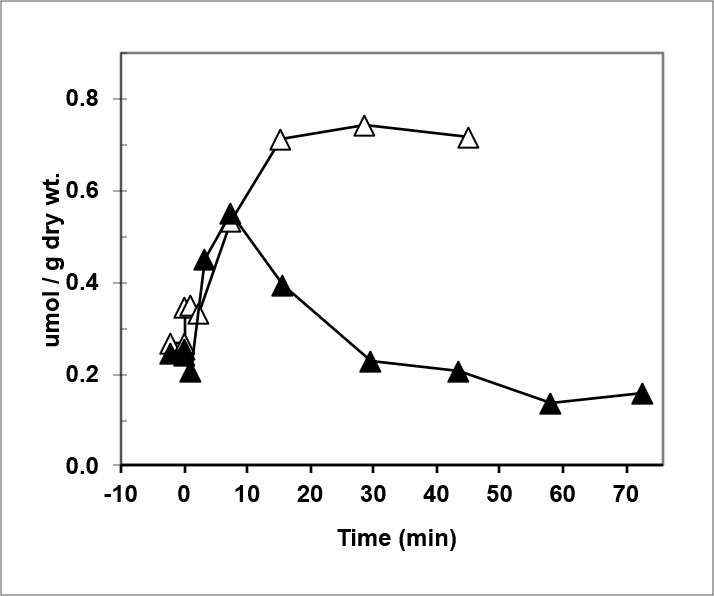
Fig. S1 Mild upshift

**dGTP**

**dATP**

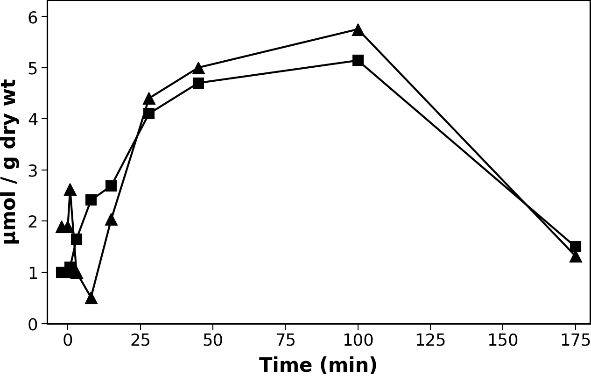

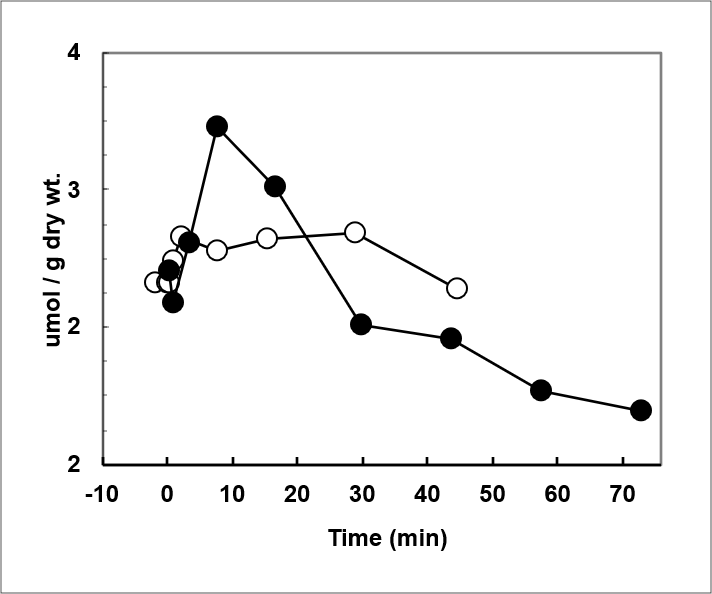

**CTP**

P

**UTP**

**GTP**

FIG S1 **Nucleotide pools following mild osmotic upshift.** Cells were subjected to mild osmotic upshift (0.45 M NaCl) at t = 0. Intracellular nucleotide levels were determined over time by radiolabeling and thin-layer chromatography. Closed symbols excess K⁺ (5mM); open symbols limiting K⁺ (0.3 mM).

Fig. S2 Moderate upshift

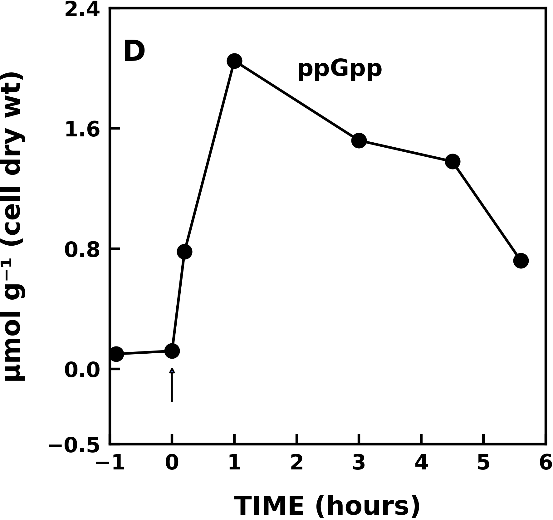

FIG S2 **ppGpp levels following moderate osmotic upshift.** Cells were subjected to moderate osmotic upshift (0.57 M NaCl) at t = 0. Intracellular nucleotide levels were determined over time by radiolabeling and thin-layer chromatography.
